# Bioinspired Dual-Functional Solid Lipid Nanoformulations for Targeted Drug Delivery and Sustained Release for Enhancement of Potency of Albendazole, an Antihelminthic Drug

**DOI:** 10.1101/2021.07.24.453620

**Authors:** Sunidhi Sharma, Vanshita Goel, Pawandeep Kaur, Kundlik Gadhave, Neha Garg, Lachhman Das Singla, Diptiman Choudhury

**Affiliations:** School of Chemistry and Biochemistry, Thapar Institute of Engineering and Technology, Patiala, 147004, Punjab, India; Indian Institute of Technology (IIT), Mandi, Mandi-175005, Himachal Pradesh-India; Institute of Medical Sciences, Banaras Hindu University, Varanasi, Uttar Pradesh 221005, India; Guru Angad Dev Veterinary and Animal Sciences University (GADVASU), Ludhiana, 141001, Punjab, India; Thapar Institute of Engineering and Technology-Virginia Tech Centre of Excellence for Emerging Materials, Thapar Institute of Engineering and Technology, Patiala-147004 Punjab, India; Vedanga Life Technologies, Thapar Institute of Engineering and Technology, Patiala-147004, Punjab, India

**Keywords:** Solid Lipid Nanoparticles, Albendazole, Sustained release, Anti helminthic drug, Targeted delivery, Pharmacokinetics

## Abstract

Targeted delivery has not been achieved for anthelmintic treatment, resulting in the requirement of excess drugs dose leading to side effects and therapeutic resistance. Gastrointestinal helminths ingest lipid droplets from digestive fluid for energy production, development, and defense. Worm’s habit inspired us to develop biocompatible, oral administrable, bee-wax derived solid lipid nanoparticles (SLN) with excellent drug (albendazole) loading efficiency of 83.3 ± 6.5 mg/g and sustained-release properties (86.4 ± 3.9 % of drug released within 24 h). Rhodamine B-loaded SLN showed time-dependent release and distribution of dye *in vivo in Haemonchous contortus*. The intestinal sustained-release property was shown by the particles that caused enhancement of albendazole potency for up to 50 folds. Therefore, this formulation has immense potential as an anthelminthic drug delivery vehicle that will not only be able to reduce the dose but will also reduce the drug-induced side effects by enhancing the bioavailability of the drug.

**Highlights:**

- Albendazole-loaded Solid Lipid Nanoparticles (SLN-A) were formulated using Beeswax as stating material showing high drug loading capacity of 83 mg/g with sustained-release properties and 84 ± 3 % of drug release within 24 h.
- SLN-A particles showed 50 fold enhancement of Albendazole activity against *Haemonchous contortus* worm.
- Rhodamine B-loaded SLN particles showed the specific uptake and *in-vivo* sustained release of dye in the worm.

## 1. INTRODUCTION

Of all the infectious pathological diseases present in humans and livestock, helminthic infection is one of the most common types. 20% of the world’s human population, almost all wild animals, and a significant portion of the farm animal in developing countries are infected with intestinal helminths. Even though the mortality rates in these infections are less, it has been shown to cause various health complications like diarrhea, malabsorption, blood loss, and reduced growth rates (Awasthi et al., 2003; “WHO | Prevention and control of intestinal parasitic infections: WHO Technical Report Series N° 749,” 2016). In some cases, they can cause anemia and retarded growth in children, adversely affecting cognition and educational abilities. In addition, the economic loss due to these soil-transmitted nematode infections results in a reduction of milk, dairy products, wool, meat, etc. (Flach, 2008; Weber, 2015). Of these intestinal worms, *Haemonchus contortus* is a type of blood-feeding nematode belonging to the Trichostrongyloidea family. It is present in small ruminants and causes haemonchosis and severe damage in the mucous membrane of the abomasum, resulting in gastric hemorrhage (Gasser and Samson-Himmelstjerna, 2016; Gelberg, 2017; Sendow, 2003). Although this worm is predominant in the tropical and subtropical regions and infects sheep and goats, a few cases of human infection were reported in human populations of Australia and Brazil (Sendow, 2003). *H. contortus* is known to absorb lipids from the GI tract of the host that is used for energy production, synthesizing parasite-specific glycerolipids and phospholipids, which are involved in host-parasitic interactions (Wang et al., 2020). In addition, a large amount of triacylglycerol was found in parasitic eggs which are used as a direct energy supply for egg development and are actively accumulated in the form of fats in the embryos (Wang et al., 2020). The uptake of lipids by the *H. contortus* worm from the host inspired us to make drug-loaded lipid nanoformulation for targeted delivery that will help in reducing the drug dosage and their side effects. Due to the presence of lipid core, the worms will specifically uptake the particles, taking the drug alongside and thereby increasing its potency.

A limited number of drugs, belonging to the macrolides and benzimidazoles family, are known to act against these intestinal worms. Albendazole (Abz) is a type of benzimidazole drug and was developed in 1975. It has been used to control a wide variety of parasitic nematodes in humans and animals for decades (Barrère et al., 2012). Due to its poor water solubility, the Abz drug is usually administered along with a fatty meal (García et al., 2014). The mechanism of action involves its binding to β-tubulin at the colchicine’s-binding site, causing inhibition of the polymerization leading to the parasite’s death (Jordan and Wilson, 2004; Kumar et al., 2013). But in recent days, chemoresistance due to altering tubulin isotype has been reported on several occasions for human and animal parasitic worms (Barrère et al., 2012; Sallé et al., 2019). Furthermore, the recommended dose of Abz against *H. contortus* is 10 mg/kg and 7.5 mg/kg of body weight for humans and cattle, respectively according to FDA (“CFR - Code of Federal Regulations Title 21,” 2020; Goel et al., 2020). However, the use of such a high dosage leads to side effects like headache, nausea, vomiting, stomach pain, dizziness, fever, loss of liver function, etc. (Samuel et al., 2014).

Different types of nanocarrier systems like carbohydrate-based, protein-based, or lipid-based carriers are being used, of which lipid-based drug delivery is recent and is known to have the advantage of better encapsulation efficiency, better solubility, and low toxicity (Jannin et al., 2008). Biocompatible lipids like stearic acid, cetyl palmitate, cetyl alcohol, trimyristin, glyceryl monostearate, etc. have been used for the formation of SLNs (Dastidar et al., 2019; Dolatabadi et al., 2015; Wissing et al., 2004). In the present study, beeswax was used as a lipid source, which is a non-toxic natural wax known for its effectiveness against ringworm, jock itch, and fungal skin infections (NS, 2004). Beeswax constitutes-four unsaturated acids (oleic, linoleic, palmitoleic, and linolenic), two saturated fatty acids (tetracosanoic and palmitic), some other hydrocarbons, and wax esters. In this, free fatty acids account for 12%–14% with a chain length of C24–C32 and free fatty alcohols (1%) have C28–C35. Linear wax monoesters and hydroxymonoesters are 35%–45% with C40– C48 and are derived from palmitic, 15-hydroxypalmitic, and oleic acids. Complex wax esters are 15%–27% and contain 15-hydroxypalmitic acid or diols linked to other fatty-acid molecules through their hydroxyl groups (Fratini et al., 2016).

Ethanol extract of beeswax was prepared using soxhlet apparatus that helped in the extraction of all the free fatty acids (Buchwald et al., 2009). This was later used for Abz-loaded Solid Lipid Nanoparticles (SLN-A) formulation tested against the intestinal parasitic worm *H. contortus*. These particles are water-soluble, showed sustained drug release, are worm targeted, show increased bioavailability of the drug inside the parasite, and thereby enhance the drug potency. Abz-loaded SLNs were made biocompatible using Poloxamer 407 coating on them. Further, the effectiveness of the formulation was compared with the FDA-approved oral Abz solution for its activity. To confirm the uptake of drug-loaded SLN particles, Rhodamine B-loaded nanoparticles were prepared and observed for the route of ingestion and diffusion of the drug inside the worm body.

## 2. MATERIALS AND METHODS

### 2.1 Materials

Rhodamine B, DMSO, Ethanol, Chloroform, Hydrochloric acid, Sodium hydroxide, Sodium bicarbonate, etc., were purchased from Loba Chemie India. Unrefined Beeswax was purchased from Arora Honey Bee Farm, Amritsar, India. Poloxamer 407, MTT, RPMI Media, and DMEM Media were purchased from HiMedia Lab, India. Hek293 cells were purchased from ATCC and maintained in the lab in a low passage number for the research work.

### 2.2 Synthesis of SLN, SLN-A, and SLN-Rh particles

Abz-loaded solid lipid nanoparticles (SLN-A) were fabricated using the double emulsion technique. The ethanolic extract of beeswax was made using the soxhlet apparatus at 78 °C. These extracts were then dried overnight in a hot air oven at 37 °C. The dried extract was later dissolved in chloroform (50 mg/ml) to make a lipid emulsion. Abz (100 mg/ml) was mixed in this lipid emulsion, which was later mixed in a 2% aqueous Poloxamer 407 solution (w/v). Finally, the solvent was Rota-evaporated to get SLN-A particles in powder form (**Fig. 1**).

**Fig. 1.**
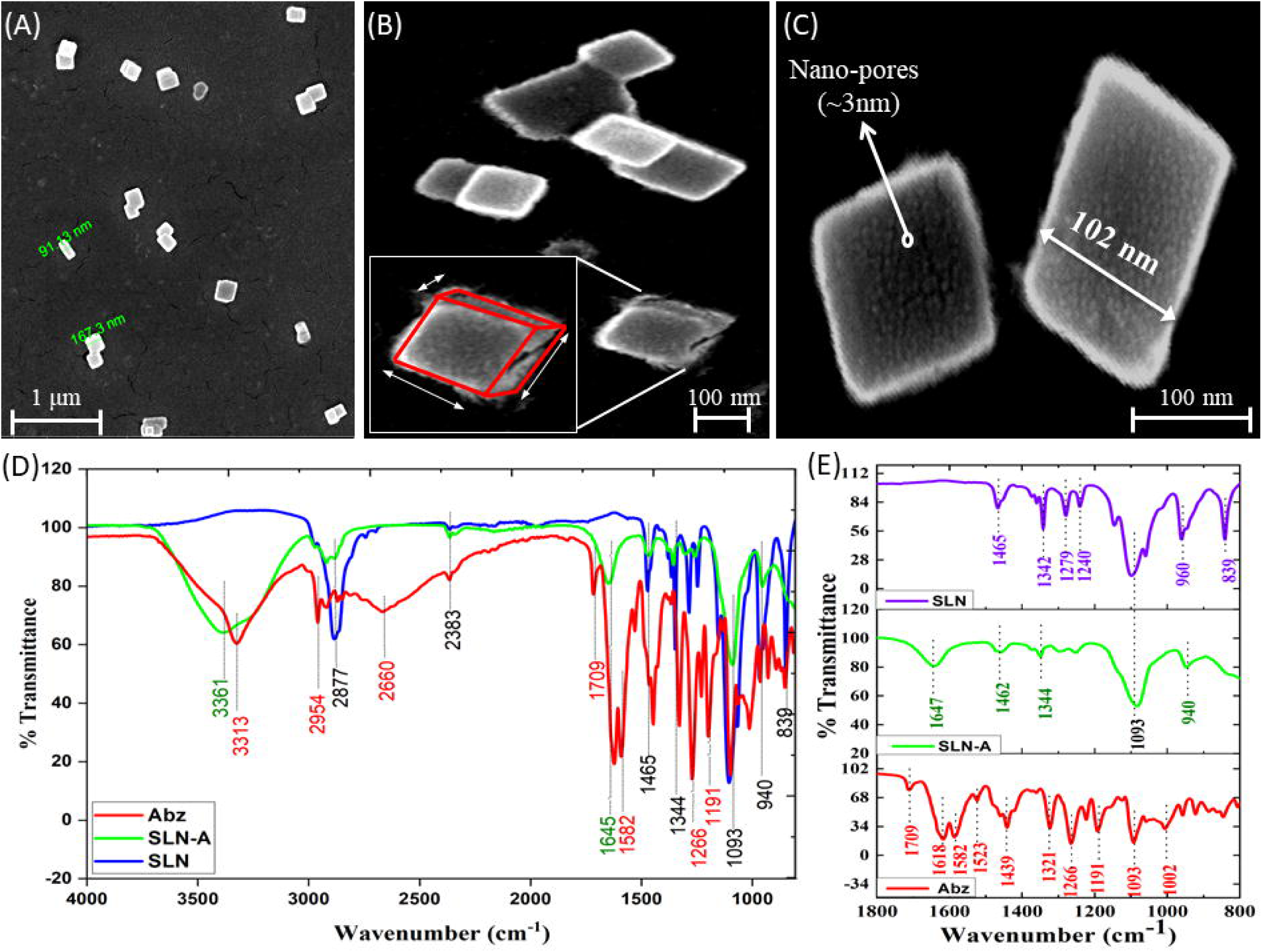
Synthesis scheme of SLN, SLN-A, and SLN-Rh particles using Beeswax and Poloxamer-407.

For the preparation of SLN particles (without drug), the particles were prepared using the above procedure without loading any drug. In addition to that, Rhodamine B-loaded SLNs (SLN-Rh) particles were also prepared using the same method. Rhodamine B (20 mg/ml) was mixed in the lipid emulsion for the formation of particles.

### 2.3 Particle size and Morphological Analysis

The hydrodynamic size of SLN and SLN-A were determined using Dynamic Light Scattering (DLS, Nanobook 90 plus, Brookhaven, USA) after dispersing the particles in distilled water. Further, the morphological characterization of SLN-A particles was examined at room temperature using the drop-casting method on a Field Emission Scanning Electron Microscope (FE-SEM, Nova Nano SEM-450, JFEI Company of USA (S.E.A.) PTE LTD). Dried samples were gold coated for 20 min before imaging. Fourier Transform Infrared Spectroscopy (FTIR, Agilent Technologies Cary 800 Series, Australia) was employed to check and confirm the drug-loading and formation of particles where the sample was scanned in powder form in the range 4000-800 cm^-1^.

### 2.4 Drug Loading and Release Kinetics

Abz drug was encapsulated in SLN-A particles and its drug loading capacity was calculated and the release kinetics was observed for SLN-A and SLN-Rh particles. The particles (1 mg/ml) were taken in 30 ml PBS buffer (pH 7.4) where the system was maintained at 37 □, and a spin of 130-150 rpm was applied. Samples were collected at regular intervals and checked for their absorbance at 295 nm for SLN-A particles and 555 nm for SLN-Rh particles using a UV-Visible spectrophotometer (UV-2600, UV-VIS Spectrophotometer, Shimadzu). Later, the percentage drug release and per hour percentage drug release were calculated and plotted against time and noted for its trend.

### 2.5 Biocompatibility of SLN /SLN-A/SLN-Rh with human cells

Albendazole and different drug/dye-loaded SLN particles (SLN, SLN-A, and SLN-Rh) were checked for their cytotoxicity using an MTT assay. Hek293 (Human Embryonic Kidney cell line) was treated with different concentrations of 5, 50, and 100 μM and incubated for 24 h in a 5 % CO_2_ incubator at 37 □ and humidified conditions. 10 μl of MTT was added and incubated for 3 h. DMSO was later added to dissolve the Formazan crystals formed, and its OD was taken using an Elisa plate reader (PowerWave XS2, BioTek) at 570 nm.

The percentage of viable cells was calculated using:

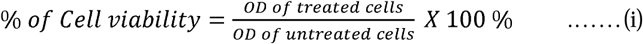

### 2.6 Isolation of adult Haemonchus contortus *worm*

Adult *H. contortus* worms were identified and isolated from the abomasum samples of goats that were collected and brought from the slaughterhouse. These worms were first washed using a 1X PBS buffer of pH 7.4 and then later transferred to petri-plates containing RPMI media. Identification of these worms was done using the compound microscope and for further experiments, worms were selected randomly.

### 2.7 Treatment with SLN-Rh particles

Randomly selected 10 worms were added to RPMI media taken in 35 mm petri-plates and treated with SLN-Rh particles (20 μM). Later the worms were incubated with SLN-Rh particles for 1 h, 2 h, 3 h, and 24 h, respectively. After the incubation, the worms were washed in 1X PBS buffer and placed on a glass slide to observe the distribution of Rhodamine B. The images were taken under the Dewinter Fluorescent Microscope at 10X and 20X magnification.

### 2.8 Treatment of worms with SLN-A particles

Worms were taken in petri-dishes containing RPMI media. Adult Motility and Morbility Assay (AMMA) was performed in which the death and paralysis time after the treatment with particles was noted. 30 worms were taken in triplets for each dose, with 15 males and 15 females in each set. SLN-A particles were added in the following concentrations-2 μM, 5 μM, 10 μM, 20 μM, 50 μM, 100 μM, and 200 μM and the Abz was taken as a standard control having concentrations - 50 μM, 200 μM, 1 mM, and 2 mM. The worms were then observed at different time intervals by checking them under the compound microscope for movement. When no movement in the worm was observed (paralyzed), the worm was touched with a drop of warm (55 °C) saline solution. If the worm doesn’t show any movement, the worm was noted as dead.

For analysis of Oxidative Stress production, 6 male and 6 female worms were randomly picked and placed in a petri-dish containing RPMI media. SLN-A particles (10 μM) were added from the stock solution. The Abz drug and SLN particles were also taken as a control using the same concentration. After incubation for 3 h, the worms were washed with PBS buffer and then incubated with DCFDA dye (10 μM) for 10 min in the dark. Later then the worms were observed under Dewinter Fluorescent Microscope at 100X and 200X magnification.

### 2.9 Statistical analysis

Data were presented as the mean of at least three independent experiments. Statistical analysis of data was conducted by a Student’s t-test, by using MS Excel, and two measurements were statistically significant if the corresponding p-value was < 0.01.

## 3. Results

### 3.1 Structure and Morphology of SLN

The SLN and drug/dye-loaded SLN particles were formed using the double emulsion technique. The flakes of SLN particles were lime-colored, orange-colored for SLN-A, and reddish for SLN-Rh particles (**Fig. 1**). These particles were found to be stable at room temperature in dry conditions. DLS data showed that the flakes of SLN-A and SLN were in the range 257 ± 98 nm and 165 ± 103 nm, respectively (**Supplementary Information, S1. A-B**). Further, the morphology of the SLN particles was confirmed using FE-SEM. The FE-SEM images showed that SLN-A particles had cuboidal-shape and their sizes range 93 ± 29 nm **(Fig. 2A-B)**. The images also showed that these particles were porous and had a pore size of about 3 nm (**Fig. 2 C**).

**Fig. 2.**
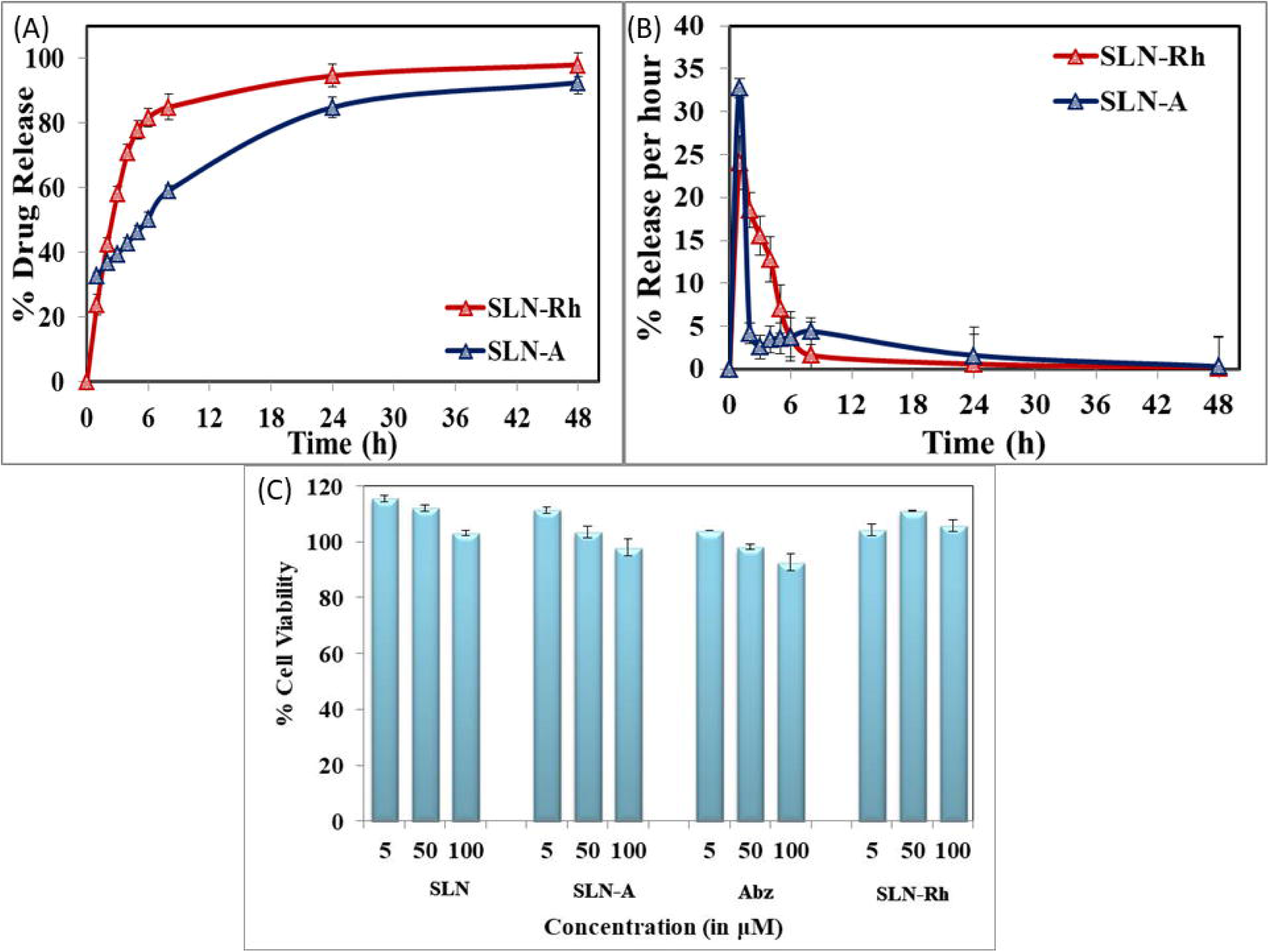
Physical characterization of SLN particles using FE-SEM and FT-IR. (A) The morphology of SLN-A particles was studied using FE-SEM image; (B) Particles showing its cuboidal shape (C) Size of these particles ranges 100 – 200 nm and have nano-pores of diameter 3 nm; (D) The FT-IR spectrum of Abz, SLN-A, and SLN particles measured using air-dried samples in the range of 500-4000 cm^-1^ using Agilent Technologies Cary 800 Series FTIR Spectrophotometer at room temperature and (E) FT-IR spectrum of Abz, SLN-A and SLN particles in the fingerprint region.

### 3.2 FT-IR study for confirming Abz loading

The FT-IR spectra were generated to determine the changes in SLN particles after the drug loading, which was clearly shown by the presence of characteristic peaks of Abz and SLN particles in SLN-A particles, proving that the Abz drug has been loaded into the particles. FTIR spectra showed a broad peak in SLN-A particles, and Abz molecule at 3313 cm^-1^ and a shift in the peak of SLN-A particles caused due to the N-H stretching at 3361 cm^-1^ from the N-H stretching at 3313 cm^-1^ in the Abz molecule. This peak was not observed in SLN particles. Another characteristic peak of the benzimidazole molecule was found in both particles due to the C=N stretching where Abz showed peaks at 1618 cm^-1^, and SLN-A showed the peak at 1647 cm^-1^ (**Fig. 2D-E**). Another peak at 2877 cm^-1^ was observed in SLN and SLN-A particles due to C-H stretching. The peak for C-O stretching was observed at 1279 cm^-1^ in SLN particles, at 1268 cm^-1^ in SLN-A particles, and at 1266 cm^-1^ in Abz molecules and C-O asymmetric stretching was observed at 1085 cm^-1^, 1073 cm^-1^, and 1182 cm^-1^ in SLN, SLN-A, and Abz respectively (Abidi et al., 2018; Gunasekaran et al., 2009; Jadhav et al., 2019). The fingerprint region showed almost similar peaks in SLN and SLN-A particles which proves the entrapment of Abz molecule inside the SLN particles in SLN-A particles (**Supplementary Information, Table S1**).

### 3.3 Drug Loading and Release Kinetics

The drug loading capacity and release kinetics of SLN-A and SLN-Rh particles were studied where the drug was loaded as 83.3 ± 6.5 mg/g in SLN-A particles, and Rhodamine B was loaded as 33.3 ± 4.2 mg/g in SLN-Rh particles. The release kinetics of both Abz and Rhodamine-B was studied from SLN-A and SLN-Rh particles at the physiological pH of 6.4 and physiological temperature of 37 □. Sustained drug release was observed in SLN-A particles for 24 h, and burst release followed by sustained drug release was observed for the first 12 h in SLN-Rh particles. Most of the drug was released by 48 h. SLN-A particles were suspended in a dialysis membrane with a concentration of 1 mg/ml, and the OD value was observed at 295 nm after particular time intervals (**Supplementary Information, Fig. S1. C**). It was observed that 86.4 ± 3.9 % of the drug was released from SLN-A particles by the end of 48 h.

On the other hand, in the case of SLN-Rh particles, OD value was observed at 555 nm after particular time intervals, and it was noted that up to 50 % of the Rhodamine B dye was released in 2.5 h, and the almost total drug was released by 24 h (93.5 ± 2.7 %) (**Fig. 3A**). This difference was observed due to the presence of polarity difference of Abz molecules and Rhodamine B molecules. Abz, due to its hydrophobic nature, is practically water-insoluble. On the other hand, in SLN-A particles, Abz is inside the lipid layer surrounded by a poloxamer molecule with its hydrophilic tail on its outer surface. Due to this, SLN-A particles have shown sustained release of the Abz drug. On the contrary, in the case of SLN-Rh, Rhodamine B is water-soluble due to its hydrophilic nature and hence has shown burst release in the first 6 h followed by sustained release of the dye molecule (**Fig. 3B**).

**Fig. 3.**
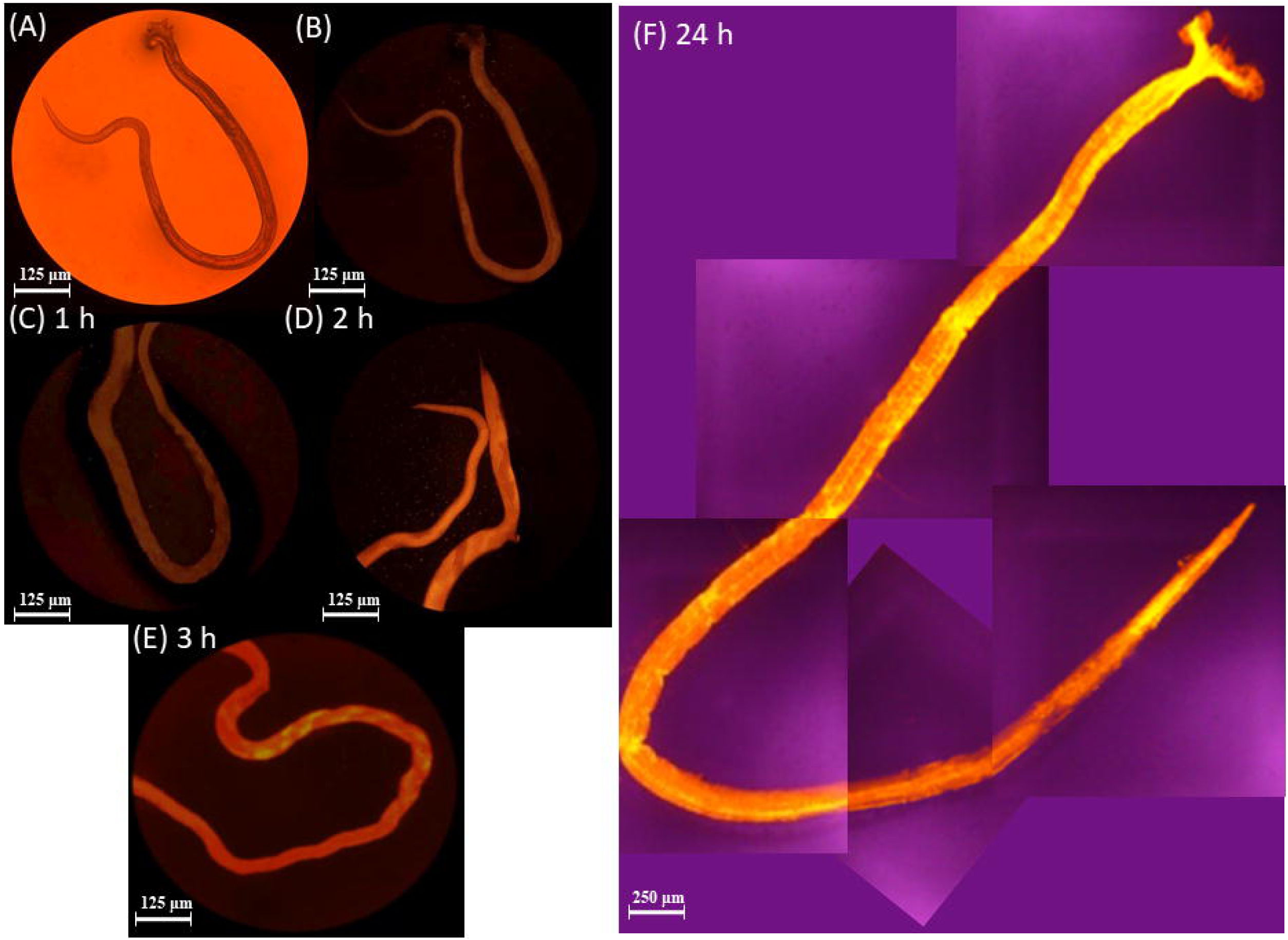
Plots showing Release Kinetics from SLN-A and SLN-Rh particles and their biocompatibility with normal cells. (A) Plot for the release kinetics of SLN-A and SLN-Rh particles showing percentage drug released from the particles; (B) Plot for the release kinetics showing per hour drug released (%) from the SLN-A and SLN-Rh particles; (C) Cytotoxicity assay of SLN, SLN-A, Abz, and SLN-Rh showing no effect on Hek293 cells.

### 3.4 Biocompatibility of SLN particles

The biocompatibility of these SLN particles was studied using the cell viability assay, i.e., MTT assay. Hek293 cells were treated with SLN and drug/dye-loaded SLN particles and checked for their viability. SLN, SLN-A particles, Abz drug, and SLN-Rh particles were treated at different concentrations of 5, 50, and 100 μM (**Fig. 3C**). No toxicity was observed for the SLN, SLN-Rh, SLN-A particles, and Abz drug compared to control untreated cells. Cell viability was found to be 103.0 ± 6.4 % for SLN particles, 97.8 ± 6.7 % for SLN-A particles, 98.1 ± 5.7 % for Abz alone, and 104.2 ± 3.5 % for SLN-Rh particles (Using equation (i). Thus, these particles show no toxicity towards normal cell lines, and they can be used as a potential drug delivery system towards parasitic helminths.

### 3.5 Treatment with SLN-Rh particles

*H. contortus* worms were treated with SLN-Rh and checked for their uptake and distribution of dye with respect to time in *H. contortus* worms. These worms were treated with 20 μM concentration for 1 h. Then images were taken in the red light filter after different time slots - 1, 2, 3, and 24 h (**Fig. 4**). An increase in fluorescence intensity was detected in worms with increasing time showing slow drug release inside the worm. Images were also taken in a blue light filter at 3 h, which showed how the fluorescence was observed more inside the alimentary canal, which was not so clear in images taken in the red light filter due to bright intensity (**Supplementary Information, Fig. S2**). Fluorescence initially observed in the alimentary canal brightened with time and was later diffused into the whole body, as seen in 24 h images. The images show that the worm first took up the particles, then the drug was released inside the alimentary canal, and later the drug was diffused in the worm body. Treatment of worm with SLN-Rh particles showed the route of particles and dye inside the body with increasing time.

**Fig. 4.**
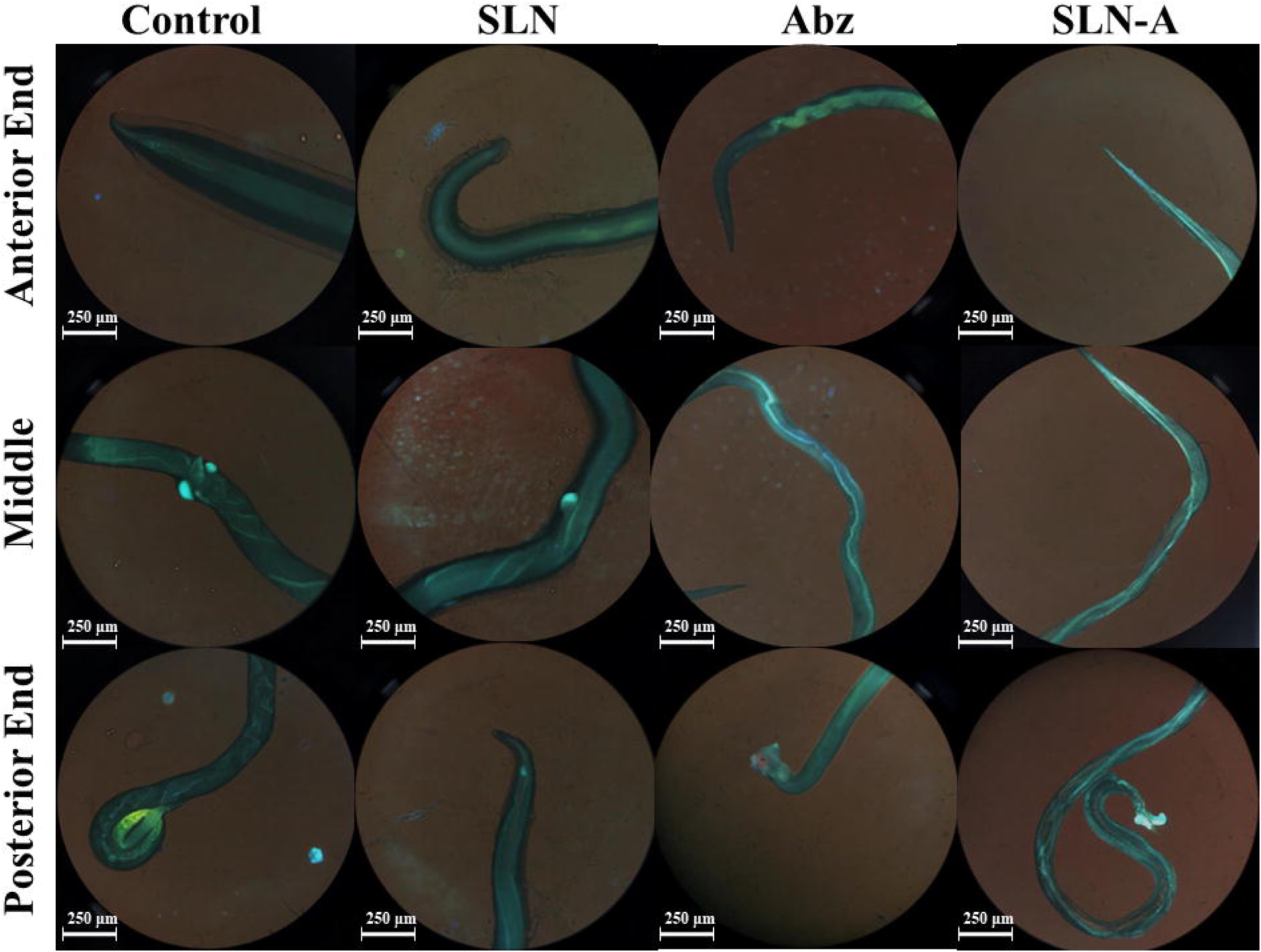
Treatment of worms with Rhodamine-B loaded SLN particles. *H. contortus* worms were treated with SLN-Rh particles and the release pattern of the dye was observed at 100X magnification in Dewinter Inverted Fluorescence Microscope, Italy. (A) Full adult worm in bright white light at 40X magnification; (B) Full adult worm after excitation of green light at 40X magnification after 24 h; (C), (D), and (E) Intestine of adult worm after 1 h, 2 h, and 3 h respectively showing increased fluorescence intensity with each passing hour; (F) Worm at 100X magnification showing diffused dye inside the whole body of worm after 24 h. This shows that the particles were ingested by the worm and the drug/dye was released which was later diffused inside the whole body.

### 3.6 Anti-helminthic activity of SLN-A particles

Adult Motility and Morbility Assay (AMMA) was performed for testing the antihelminthic activity of SLN-A particles, where the death and paralysis time of *H. contortus* worms were noted upon its treatment. 15 adult males and 15 adult females each were utilized in triplets, and it was observed that at 10 μM concentration, 66 % of the worms were paralyzed in 12 h and 100% of worms were dead within 24 h of treatment at 20 μM SLN-A particles. As a control Saline, PBS (pH 7.4) and RPMI media were used, and no deaths were observed for 24 h. In the case of Abz alone, worms had started showing paralysis at 6 h, and deaths were observed after 12 h for the 1 mM and 2 mM concentrations, and no death was observed for the lower concentrations till 24 hours. While in the case of SLN-A, lower concentrations like 5 μM showed paralysis at 6 h and death at 12 h. After 24 h, almost all the worms were found dead, showing better outcomes than in Abz alone (**Table 1**). For adult male and female worms, the LD_50_ values were 1 mM and 10 μM for Abz and SLN-A particles treatment.

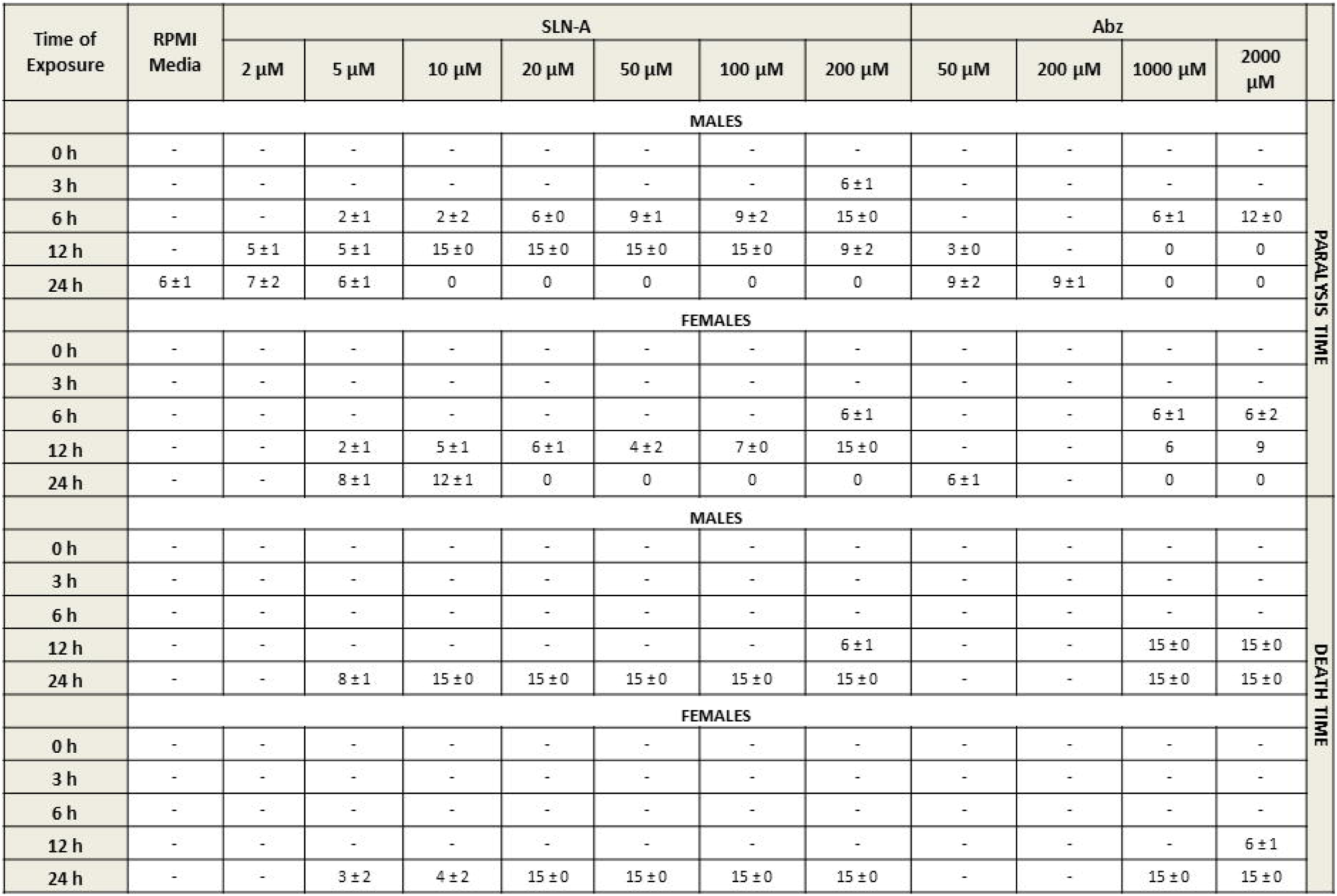

### 3.7 Reactive Oxygen Species (ROS) stress generation due to SLN-A treatment

Oxidative stress generation in the adult worms due to the treatment of Abz, SLN, and SLN-A particles was checked using ROS assay (**Fig. 5**). After the incubation with particles, the worms were stained with DCFDA dye and the images were taken in the violet light filter and checked for ROS generation. The images were taken at the anterior, middle, and posterior end of the adult worms, and the difference in intensity of fluorescence showed the amount of stress generated. In SLN-A and Abz treated worms, high intensity of green fluorescence was observed, showing greater production of oxidative stress and more damage to the tissues. Among these, SLN-A particles showed greater ROS production with pixel intensity of 113.2 × 10^4^ ± 36.3 when compared to the worms treated with Abz alone showing pixel intensity of 102 × 10^4^ ± 27.1 (**Supplementary Information, Fig. S3)**. On the other hand, very little ROS production was observed in Control worms (RPMI media) with pixel intensity of 43 × 10^4^ ± 8.9 and SLN treated adult worms pixel intensity of 52.9 × 10^4^ ±10.2.

**Fig. 5.**
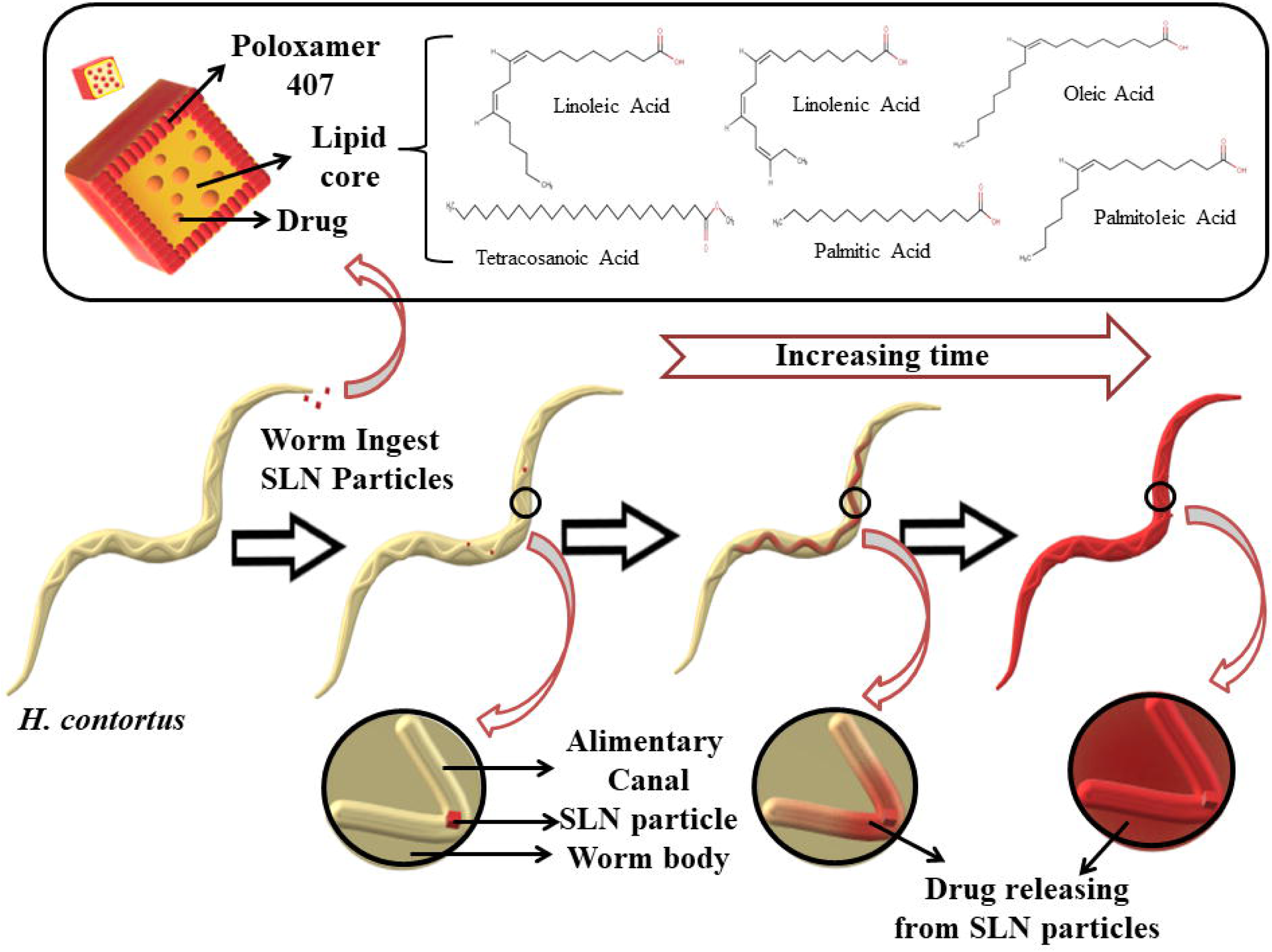
Study of oxidative stress generated in *H. contortus* adult worm. ROS assay was performed to check the amount of oxidative stress generated when the worm was treated and incubation with SLN, Abz and SLN-A particles for 6 h. The staining of worms was done using DCFDA dye (100 nM) and the amount of ROS generated was observed in the anterior, middle and posterior regions of the worm at 100X magnification in Dewinter Inverted Fluorescence Microscope, Italy.

## 4. Discussion

Benzimidazole type of drugs have been used commonly as a treatment for parasitic worms, but with increasing cases of resistance against these antihelminthic drugs, there is an urgent need for the development of a new drug or a better drug delivery carriers option. While the development of a new drug requires a lot of time and lots of resources, other drug delivery carrier systems can be formulated to increase the efficacy of already available antihelminthic drugs. *H. contortus* is a highly pathogenic intestinal parasitic worm that causes gastrointestinal infections in ruminants and sometimes in humans (Goel et al., 2021). These worms are known to uptake and utilize lipids from the hosts for egg and larval development, energy production, and synthesizing parasite-specific glycerolipids and phospholipids required for host-parasitic interactions (Wang et al., 2020). With increasing cases of antihelminthic drug resistance in the worms, drug-loaded SLN particles can prove to be of great benefit in oral drug delivery applications.

Here in this work, lipids that were extracted from beeswax were used for the formulations of Abz-loaded solid lipid nanoparticles (SLN-A). Beeswax, also known for its anti-allergenic and anti-swelling properties, have been used in pain-relieving, lowering cholesterol, reducing diaper rashes, and have an effect against ringworm and jock itch (NS, 2004). Therefore, beeswax, being the natural wax having various health benefits and a great source of lipids was used for these nanoparticle formulations. Poloxamer 407, because of its amphiphilic nature, acts as a surfactant and stabilizing agent and has been used as a co-polymer on SLN particles which made them biocompatible and increased the availability of drugs inside the system.

Taking into consideration the above factors, SLN-A particles were made that showed enhanced antihelminthic activity of Abz against the resistant parasitic worm, *H. contortus* when compared with the drug alone. SLN-A particles not only increased the aqueous solubility of Albendazole, a BCS Class-II drug but also increased its effectiveness. These particles were specifically uptaken by the adult worm, which was further supported by SLN-Rh particles where a time-dependent Rhodamine B release was observed in the intestine and then was later diffused in the whole of the adult *H. contortus* worm. Specific uptake and intestinal sustained-release dramatically enhanced the activity of SLN-A particles in the intestine for up to 50 fold. The biocompatibility of these particles was checked for their cell viability on Hek293 cells using MTT assay and was found to be non-toxic on treatment with SLN, SLN-A, and SLN-Rh particles. Hence these particles showed no side effects, and the antihelminthic activity was due to the Abz drug alone inside the SLN-A particles.

Further, Adult Motility and Morbility Assay (AMMA) was performed on the *H. contortus* worms where the effect of SLN-A particles was checked on the adult worms for their death and paralysis. The IC_50_ value for SLN-A was found to be 10 μM and in the case of Abz was found to be at 1 mM. This showed that lower drug dosage of SLN-A particles showed better efficacy when compared to Abz drug alone. Oxidative stress due to SLN-A treatment was checked using ROS assay where SLN-A particles generated more oxidative stress and greater tissue damage in the worms than Abz alone showing better potency and effectiveness. Because of reduced drug dosage for these particles and better bioavailability, drug-loaded SLN can be used as suitable carrier systems for the oral drug route as these will have reduced side effects, better drug distribution, and specific targetting.

Even though mortality rates in helminthic infection are less, these infections cause malnutrition in adults but may be fatal for newborns. Nearly 44 million pregnant women in developing countries are infected with soil-transmitted gastrointestinal worms that result in birth defects, various developmental complications of the foetus, and morbidity (Geerts and Gryseels, 2000; Salawu et al., 2020). Antihelmintic drugs like Albendazole, Mebendazole, and Ivermectin do not show targeted delivery and have a high drug dosage of 200-400 mg/day, which has lead to therapeutic resistance in different gastrointestinal parasitic worms like roundworms, whipworms, hookworms, flukes, etc. and many side effects in humans and animals (Geerts and Gryseels, 2001, 2000). Therefore, albendazole-loaded SLN formulations, which are target-specific and require much less drug dosage, can be utilized for other parasitic worms as well which will improve the effectiveness of the already existing drugs.

## Acknowledgments

S.S. would like to thank TIET, V.G and L.D.S for support and giving acceptance to carry out animal experiments. V.G. would like to acknowledge TIET for providing fellowship. P.K. would like to acknowledge DST-Inspire. D.C. would like to thank the Thapar Institute of Engineering and Technology-Virginia Tech (USA) Centre for Excellence and Material Sciences for providing funds. L.D.S. is willing to thanks ICAR and GADVASU. N.G. acknowledges IIT Mandi (BioX center and AMRC) for providing experimental facilities and is grateful to Ramanujan SERB grant, India (SB/S2/RJN-072/2015).

## Figure Legends

### Graphical Abstract

SLN particles containing Drug/Dye and lipids like Linoleic acid, Linolenic acid, Oleic acid, Tetracosanoic acid, Palmitic acid and Palmitoleic acid, obtained from beeswax were encapsulated inside Poloxamer 407. These SLN particles were ingested by the *Haemonchus contortus* worms which were ingested through oral route and drug release was observed in the alimentary canal and then later diffused inside the whole adult worm with increasing time.

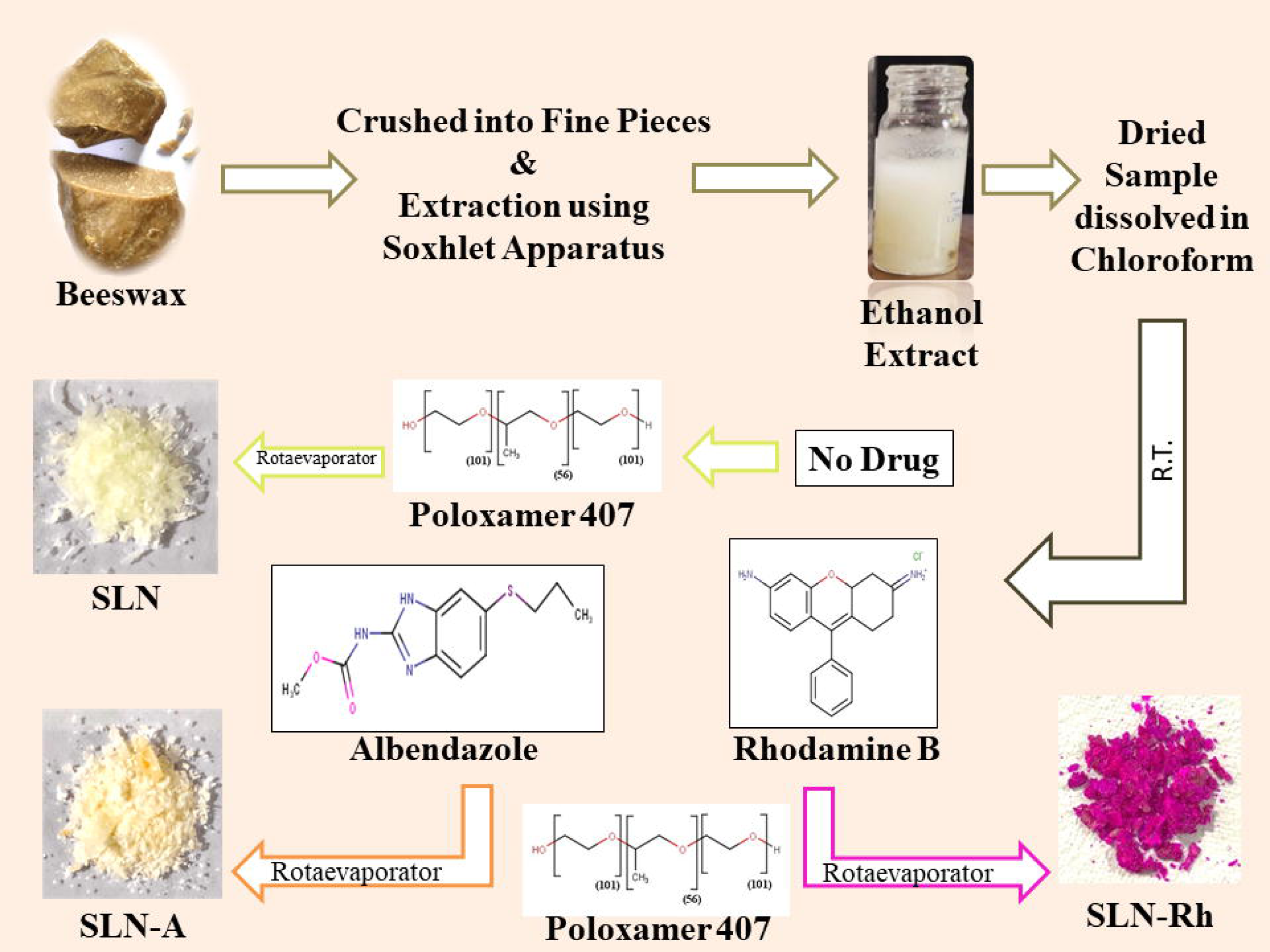

