## Supplementary material for "Bioinspired Dual-Functional Solid Lipid Nanoformulations for Targeted Drug Delivery and Sustained Release for Enhancement of Potency of Albendazole, an Antihelminthic Drug": Supplemantary Data

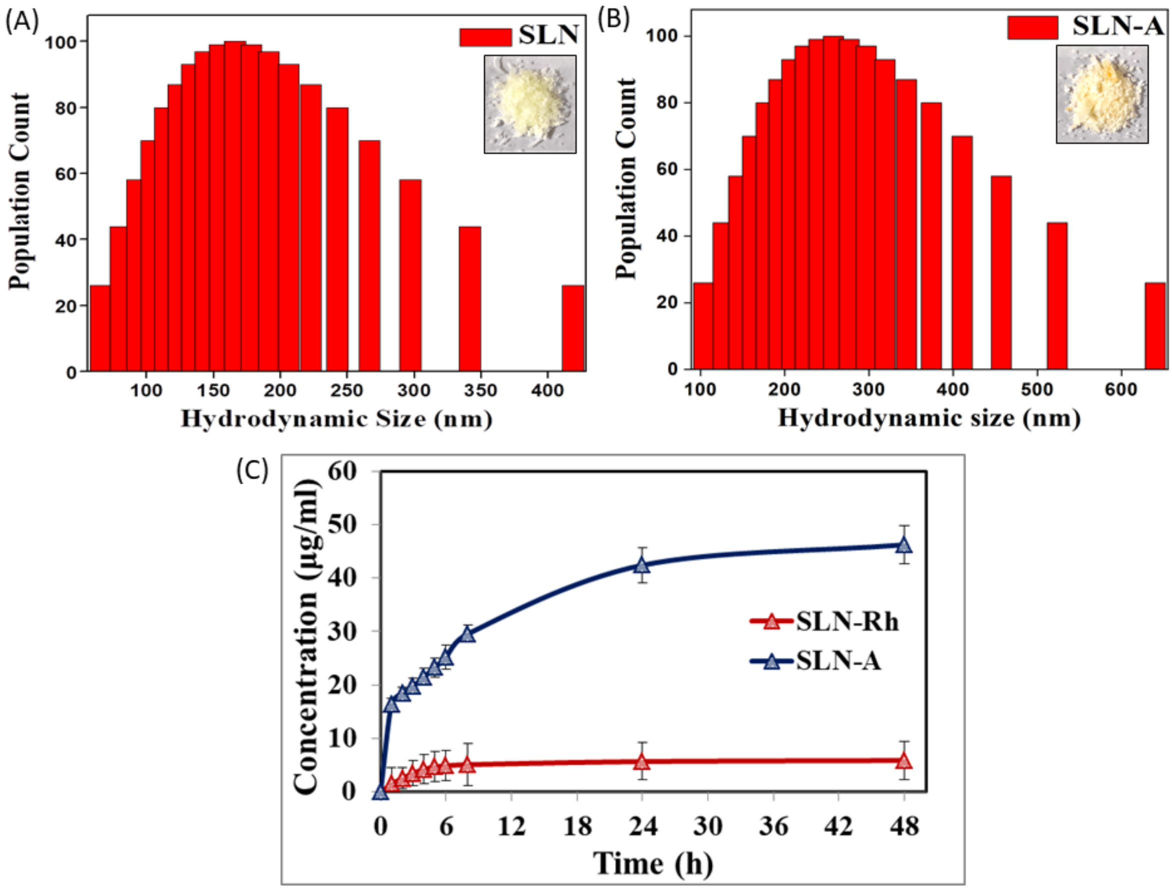


**Fig. S1. Physical characterization and Drug release pattern of SLN particles.** (A) and (B) Particle size of SLN and SLN-A particles was measured using DLS (Inset: SLN and SLN-A particles); (C) Plot showing the amount of drug released (in µg/ml) from SLN-A and SLN-Rh particles.

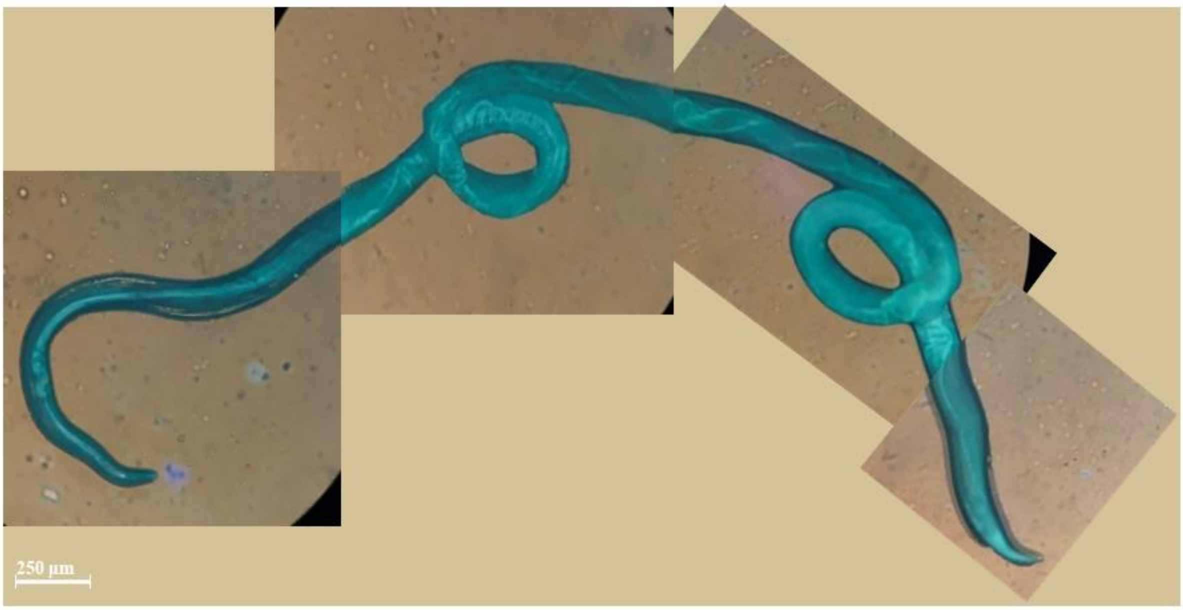


**Fig. S2. Treatment of Rhodamine-B loaded SLN particles.** *H. contortus* worm were treated with SLN-Rh particles and incubated for 3 h and then looked for fluorescence in Dewinter Inverted Fluorescence Microscope, Italy in blue filter light at 100X magnification. This filter showed greater fluorescence inside the gastrointestinal tract when compared to outside body proving that the particles were ingested by the worm and the drug/dye was released which was later diffused inside the whole body.

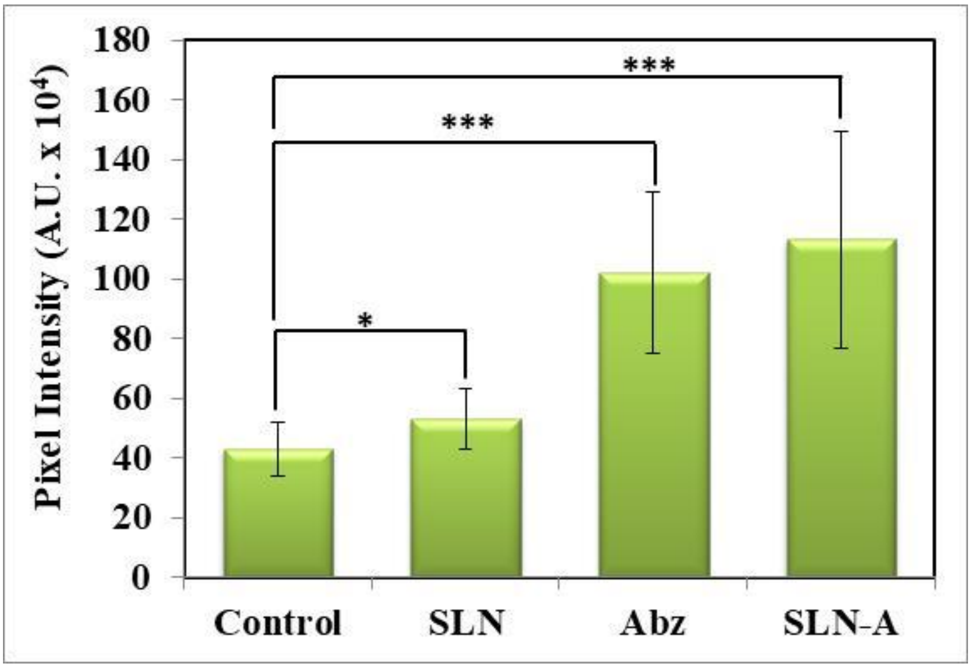


**Fig. S3.** Pixel intensity plot of DCFDA fluorescence after performing ROS assay with SLN, SLN-A, and Abz particles on *H. contortus* worm (***p-value < 0.001, ** p-value < 0.05, * p-value < 0.01).

**Table S1.** The FT-IR spectrum was measured using air-dried samples and scanned in the range of 500-4000 $cm^{-1}$.

| Remarks | SLN | SLN-A | Abz |
|---|---|---|---|
| N-H Stretching | - | 3361 | 3313 |
| C-H Stretching | 2877 | 2877 | 2954 |
| unknown | - | - | 2660 |
| C=N Stretching | - | 1647 | 1618 |
| C-H Scissoring | 1465 | 1462 | 1439 |
| C-H Bending | 1342 | 1344 | - |
| N-H Stretching | - | - | 1321 |
| C-O Stretching | 1279<br>1240 | 1268 | 1266 |
| C-O Asymmetric Stretching | 1085 | 1073 | 1182 |
| Fingerprint Region | 1093<br>960<br>839 | 1093<br>940<br>- | 1191<br>1093<br>1002 |
